# Diverse root fungal endophytes mediate plant access to soil nutrients

**DOI:** 10.64898/2026.06.27.735019

**Authors:** Rachel A. Hammer, Marissa R. Lee, Nathaniel Yang, Megan Kan, Noah Luecke, Maggie Wilson, Rhona K. Stuart, Christine V. Hawkes

## Abstract

Plant roots are broadly colonized by endophytic fungi with saprotrophic capabilities, but our understanding of whether they function in ways that are beneficial or detrimental to the host remains limited to model organisms. We hypothesized that endophytic fungi broadly affect plant access to soil nutrients, particularly organic forms that are typically not directly available to the plant. To address this, we paired 41 fungal endophytes with switchgrass (*Panicum virgatum* L.) and provided either inorganic or organic forms of nitrogen (N) and phosphorus (P). We evaluated how the fungi affected plant tissue N and P as well as plant growth. We also examined if these outcomes could be predicted from fungal phylogenetic relationships, in vitro traits of the fungi, or characteristics of the habitat from which fungi were isolated. There was substantial variation in both plant N (0.05-0.63%) and P (0.02-0.10%) acquisition that depended on the interaction of fungus and nutrient treatment. More fungi were beneficial for plant N than for P and shoot nutrients generally increased more than root nutrients from fungal associations. However, fungal effects on plant nutrients were not predicted by fungal traits, habitat traits, or fungal phylogenetic relationships. This unpredictability highlights a key challenge for incorporating endophytes into nutrient management strategies. Improving our ability to predict endophyte impacts on host nutrient acquisition will require identifying the mechanisms underlying observed beneficial effects and scaling up to realistic, diverse root microbial communities.

## INTRODUCTION

Plants can readily acquire inorganic forms of essential nutrients, but access to organic forms typically requires microbial mediation. Although arbuscular mycorrhizal fungi are a key player in plant acquisition of phosphorus (P; Wu *et al*., 2024), plant roots are also colonized by diverse endophytic fungi, including Ascomycota (e.g., Figure 1), Basidiomycota, and Mucoromycota (Li *et al*., 2025). Historically, these root endophytic fungi were defined as inhabiting roots without causing disease and were considered to be commensals, but an increasing body of work suggests they can affect biogeochemical cycles (Wang *et al*., 2021; Netherway *et al*., 2024) and plant nutrient phenotypes (Santos *et al*., 2021).

**Figure 1.**
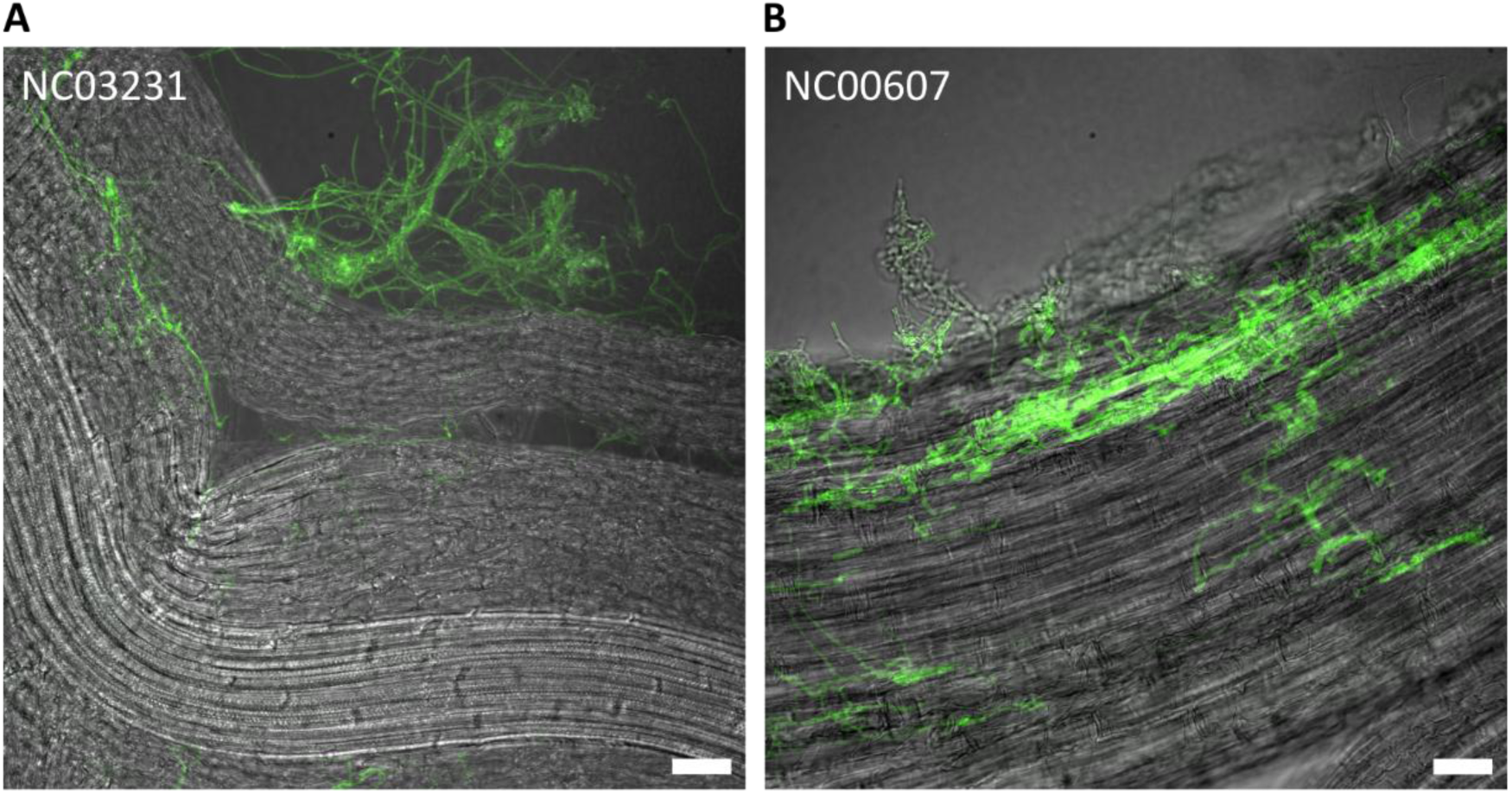
Examples of endophytic fungi colonizing roots. Images for all fungal strains are provided in Fig. S4-5. Scale bar represents 50 μm.

Endophytic fungi are horizontally transmitted from the environment, where they can also live as saprotrophs with the capacity to enzymatically degrade complex organic matter (Caldwell *et al*., 2000; Mandyam *et al*., 2010). In doing so, endophytic fungi have the potential to mobilize nutrients from otherwise inaccessible organic pools and make nutrients more available in soil (L’Espérance *et al*., 2024). Both soil saprotrophic fungi and arbuscular mycorrhizal fungi may specialize on factors such as resource quantity or quality (Johnson *et al*., 2010; Pugnaire *et al*., 2023), suggesting the potential for a range of resource-related capabilities in the endophytic fungi. This is supported by comparative genomics analysis of endophytic fungi, which reveals an expanded repertoire of enzymes, including some for degrading plant cell walls, indicating a capacity for complex substrate degradation (Knapp *et al*., 2018). Based on our knowledge from other fungi, we expect that root endophytic fungi have similar degrees of functional variation that can alter nutrient availability to plants.

Because endophytic fungi can live partly in soil and partly in the root, they also have the potential to transport nutrients to the plant akin to mycorrhizal fungi. Growing evidence suggests that nutrient transfer from endophytic fungi to the plant host may be common. For example, entomopathogenic root endophytic fungi can transfer isotopically labeled ^15^N from dead insects to their host plants, although the amount of transfer depended on both the identity of the fungus and the host (Behie & Bidochka, 2014). Also, the “mycorrhizal-like” Mucoromycotina fungi were able to transfer ^33^P and ^15^N to clover in a gnotobiotic system while receiving plant-fixed ^13^C-labeled carbon (Hoysted *et al*., 2023). Similar results were observed in traditionally non-mycorrhizal Brassicaceae species whose roots were colonized by an endophytic fungus (Almario *et al*., 2017). Other fungal strains isolated from roots transfer nutrients to the plant only under certain conditions, such as limited nutrients (Hiruma *et al*., 2016) or predominantly organic nutrient sources (Peng *et al*., 2022). Although these studies support a potential role for endophytic fungi in plant nutrient access, focusing on individual strains limits our ability to generalize more broadly or to begin predicting which fungi will be beneficial under which contexts.

Root fungal endophytes are found in diverse host plants, but are particularly common and abundant in grasses (Jumpponen & Trappe, 1998; Mandyam & Jumpponen, 2008). Switchgrass (*Panicum virgatum* L.) is a warm-season, perennial grass that is widely distributed across North America, excluding the westernmost states (Barney & DiTomaso, 2010). Because switchgrass can establish and persist in marginal soils, it is a strong biofuel candidate, but to grow this bioenergy crop with minimal inputs will rely on microbial associations for nutrient acquisition (Hestrin *et al*., 2021). Research shows that switchgrass roots are routinely colonized by endophytic fungi (Kleczewski *et al*., 2012; Singer *et al*., 2019; Dirks & Jackson, 2020; Lee & Hawkes, 2021a,b), although their role in switchgrass nutrient acquisition remains largely unknown (Hestrin *et al*., 2021).

To examine how diverse endophytic fungi affect plant access to nutrients, we grew switchgrass with 41 individual endophytes previously isolated from switchgrass roots. Plants were provided with either inorganic or organic forms of N and P, and nutrient acquisition in plant tissue was measured after 3 months. We hypothesized that root endophytic fungi would have a range of effects on plant nutrient acquisition from both inorganic and organic sources. Given their putative enzymatic capacities, we expected that the majority of endophytic fungi would have access to organic nutrients, which would make these more accessible to the plant. Inorganic nutrients are already bioavailable, and thus we expected that some fungi would successfully compete with the plant for these nutrients, resulting in a reduction in plant nutrient acquisition. While fungi have a suite of tools for accessing nutrients, we hypothesized that there may be a trade-off in ability to improve plant access to either N or P, rather than benefit the plant for both equally. Related to this, we identified fungi that were generalists or specialists for plant acquisition of N and P. We also asked whether these outcomes could be predicted from fungal traits that might drive plant-fungal interactions, including phylogenetic relatedness, enzyme activities, growth rates, and root colonization.

## METHODS

### Isolation and selection of fungal endophytes from roots

We sampled switchgrass (*Panicum virgatum* L.) across a 462-km range in North Carolina, USA, in Fall 2018 (14 sites) and Summer 2020 (repeated sampling at 4 sites). At each site and date, we collected roots from 8 plants in a stratified random design, using a shovel to dig up roots to 15-cm depth immediately adjacent to the north and south side of the plant. Soils associated with each plant were also collected for measurements of total inorganic N (TIN), P, and soil organic matter (SOM) in the form of humic matter through the NC Department of Agriculture and Consumer Services soil testing service.

Samples were stored in plastic bags on ice for transport to the lab, where fine roots < 2 mm were separated from soils and washed within 48 h. Roots were then surface sterilized in in 95% ethanol for 30 s, 3% sodium hypochlorite for 30 min, and 70% ethanol for 2 min, followed by 2 min in sterile water. After surface sterilization, roots were cut into 2-mm fragments (60 fragments per plant, 480 fragments per site, 8640 total), pressed onto potato dextrose agar (PDA) to check for residual surface contamination, and then placed into a 2-mL slant tube of PDA (1 fragment per tube). Tubes were regularly checked and fungi that emerged were transferred to PDA plates. Any fragments that showed contamination were discarded (<1%). This process resulted in 1224 isolates, of which 798 strains representative of morphotype groups were identified by Sanger sequencing of the LSU rDNA with primers NL1 and NL4 (O’Donnell, 1993) at the NC State University Genomic Sciences Laboratory. For experimentation, we selected 41 root fungi that represented both their phylogenetic diversity and range of collection environments (Table S1), excluding known pathogens and strains that were also found in plant leaves collected at the same time (Lee & Hawkes, 2021a). All fungi were also re-tested for ability to colonize switchgrass root (see Supplementary Material).

### Experimental design and measurements

To examine the role of root endophytes in host access to soil nutrients, we grew switchgrass with individual endophytes and either inorganic or organic sources of N and P. Switchgrass seeds (cv. Performer; Ernst Seeds, Meadville, PA, USA) were first surface-sterilized by soaking in 95% ethanol for 15 s, 5% NaOCl for 3 min, and 70% ethanol for 2 min, followed by 4 rinses with sterile water. Seeds were placed in 15-mL tubes of Murashige and Skoog (MS) agar with 1 drop of 500 ppm gibberellic acid and were placed in the dark at room temperature for 1 week, then moved to a growth chamber for 4 weeks with 26/24 °C day/night temperatures and 14 h days. Plants were then removed from the media for inoculation with each of the selected fungi or as fungus-free controls.

To generate the fungal treatments, we cultured each fungal strain in potato dextrose broth (PDB) with 50 µg mL^-1^ ampicillin and 10 µg mL^-1^ gentamycin for 7-10 days in 50-mL tubes. The fungal cultures were then centrifuged (3220 g for 10 min) and rinsed with sterile water three times before resuspending in water and chopping in a blender for 15 s. We quantified hyphal fragment abundance via microscopy using a hemocytometer and adjusted hyphal concentrations to 50,000 fragments mL^-1^. Seedling roots were soaked in the culture for 36 h, then transplanted to 0.4 L pots (5 cm wide × 25 cm deep; Stuewe & Sons, Tangent, Oregon, USA) filled with a 1:1 mixture of sterilized quartz sand (Play Sand, Quikrete, Atlanta, GA, USA) and lightweight expanded clay aggregates (Hydroton, Hawthorne Gardening Co., Vancouver, WA, USA).

The pots received one of two nutrient treatments, inorganic or organic, both of which contained 2.2 mM N and 0.25 mM P. The inorganic nutrient treatment included ammonium nitrate and monosodium phosphate. The organic nutrient treatment included a mixture of casein and glycine for N (each providing 50% of the N supply) and phytic acid for P. These were added to a modified Hoagland’s solution and applied three times per week over the first three weeks of growth to achieve a cumulative addition of 9.2 mg N kg^-1^ and 2.3 mg P kg^-1^. Treatments were split into two temporal batches to accommodate growth chamber space limits, with batch-specific controls. The first batch included 5 replicates, which was increased to 7 for the second batch to account for potential mortality (total n=518 pots). Growth chamber conditions were maintained as noted earlier and pot position in the growth chamber was randomized every 2 weeks.

Plant height was measured at transplant and subsequently, height, green leaves, and survival were measured at 3, 6, and 12 weeks. Height growth rate was calculated over time. Plants were harvested after 13 weeks. Aboveground leaves and stems were clipped and oven dried at 70 °C for biomass. Roots were picked from soil, washed in sterile water and immediately scanned (Epson Perfection V600 Flatbed Scanner, Epson America, Inc., Los Alamitos, CA, USA). Scanned images were later analyzed for root length with RhizoVision Explorer v.2.0.3 (Seethepalli & York, 2020) using the algorithms described in Seethepalli et al. (2021). Fresh roots were then weighed and subdivided for measurements of dried biomass (as above) and root fungal colonization. Total dry root biomass was calculated from the ratio of fresh to dry weight in the subsample. Specific root length (SRL) was calculated by dividing total root length by total dry root biomass. For fungal colonization, roots were fixed in 70% ethanol, stained with aniline blue, and percent root occupancy was quantified via the magnified intersection method (McGonigle *et al*., 1990). Dried shoot and root material were ground using a mixture of 2.3- and 3.2-mm stainless steel beads in a bead beater (MBB-96, Biospec, Bartlesville, OK, USA) and submitted to the NCSU Environmental Analysis and Testing Lab for quantification of tissue N and P. We report both shoot and root N and P, but focus on nutrients in aboveground biomass as a measure of plant nutrient acquisition to avoid the confounding of plant and fungal nutrient pools in roots (Tellenbach *et al*., 2010; Hobbie & Högberg, 2012). Nitrogen and phosphorus use efficiency (NUE and PUE) were calculated as recovery of N or P based on total plant nutrient content (percent N or P multiplied by biomass) divided by the total amount of nutrient applied during the experiment.

### Fungal traits

To help understand possible drivers of fungal effects on the host plant nutrient acquisition, we characterized a set of putatively related fungal traits, including habitat traits, growth on N and P, enzyme activities related to N and P, and their ability to colonize roots.

Habitat traits based on the site of origin of each fungus were obtained from Lee and Hawkes (Lee & Hawkes, 2021a), including total inorganic N (TIN), soil P, and organic matter measured as humic content (HM). We selected these traits because these might serve as selective pressures on the ability of the fungi to obtain inorganic and organic nutrients.

To examine fungal growth on inorganic and organic N and P in culture, we cultured the fungi on standard M9 minimal media with inorganic N and P (BD Difco, Franklin Lakes, NJ, USA) or on custom minimal media amended with organic sources of N or P. Specifically, N and P were provided as either (1) inorganic N (ammonium chloride) and P (sodium phosphate and potassium phosphate), (2) organic N (casein) with inorganic P as above, (3) organic P (phytic acid) with inorganic N as above. Each treatment included three replicates per fungus. Across all media, nutrient concentrations were maintained at 19.56 mM N and 48.1 mM P regardless of source and glucose (22.2 mM) was added as the carbon source. After 10 days, we quantified growth rates as the elliptical area over time.

We measured representative fungal enzyme activities for N (N-acetyl-β-D-glucosaminidase; NAG) and P (acid phosphatase; AP) using a fluorescent microplate assay (Saiya-Cork *et al*., 2002; German *et al*., 2011) for 38 fungi. The remaining three isolates produced responses that were too variable to be considered reliable. The standard soil procedure was modified for fungal isolates, which were grown on potato dextrose agar (PDA) with 50 µg mL^-1^ ampicillin and 10 µg mL^-1^ gentamycin for 7-14 days until enough biomass was produced. Six mycelial plugs from the leading edge of growth were collected and weighed, then blended in a sodium acetate buffer and otherwise processed as for soil samples. Outliers were removed from within-microplate replicates using a measure of spread, S_n_ (Jones, 2019), and remaining replicates were averaged.

### Statistics

All analyses were carried out in R v4.1.1 (R Core Team, 2021) and figures were made using ggplot2 v3.5.1 (Wickham, 2016).

Plant responses included biomass (shoot and root), relative height growth rate (RGR), root morphology (SRL), and shoot and root nutrient content (N and P). We standardized the plant responses as the deviation from fungus-free controls (d). The d-value was calculated by subtracting the batch-specific control means for each nutrient treatment from the experimental replicates. Positive d values indicate that treatment responses were greater than controls, while negative values indicate smaller treatment responses compared to controls.

To address the effects of the fungus and nutrient treatments, we used ANOVA to test individual plant responses (d values) as a function of fungus, nutrient treatment, and their interaction using the stats (v4.1.1, (R Core Team, 2021) and car (v3.0-12, Fox & Weisberg, 2019) packages. To account for multiple measurements made on the same plant (tissue N and P, NUE, PUE, biomass, growth rate, root morphology), we used a Bonferroni-corrected α = 0.005. For models showing significant treatment effects on plant responses, we used 95% confidence intervals to assess whether individual fungal effects within each nutrient condition significantly differed from zero. Given our focus on the role of root endophytic fungi in plant nutrient acquisition, we focus on the plant nutrient responses in shoots based on their independence from fungal tissue embedded in roots. Root nutrient responses, as well as biomass responses are presented in the Supplementary Material.

Potential trade-offs between fungal effects on host N and P were assessed with linear regression. Dependent variables were the raw shoot or root N and P percent, which were averaged by fungal treatment within each nutrient treatment to avoid pseudoreplication. For each nutrient treatment, we regressed shoot or root N percent on shoot or root P percent, respectively.

Fungal isolates were further classified based on mean effect sizes (d) for shoot N and P in the nutrient treatments. Fungi that significantly increased one nutrient or form of nutrient were classified as specialists, whereas those that increased both shoot N and P were classified as nutrient generalists. Fungi that decreased plant nutrients were further identified as competitors and isolates with no significant effects on plant nutrients were considered commensals. We focused on shoot nutrient content for this classification because root nutrients represent the combined pools of plant and fungal biomass (Pánek *et al*., 2024). We also used PCA to examine how this classification scheme related to the distribution of all plant response traits.

We used phylogenetic autocorrelation to determine whether fungal relatedness constrained fungal treatment effects on plant nutrient acquisition. The fungal phylogeny was generated with T-BAS v2.3, which places LSU sequences on a reference tree (Carbone *et al*., 2019). Taxonomic identities were inferred from reference tree placement, with multilocus placement used when ITS sequences were available to improve taxonomic resolution (Table S1). To test for phylogenetic autocorrelation of the measured plant response traits, we used Abouheif’s C_mean_ (adephylo v1.1–13; Jombart *et al*., 2010). We did the same for independent fungal traits, designed to understand whether trait analyses required a phylogenetic approach. Because we found no phylogenetic signals in either plant response traits or fungal traits (Table S2, Figure S1), we did not account for phylogenetic information in the statistical analyses.

To evaluate the relationship between fungal traits and fungal effects on plant size and nutrient acquisition, we used ordinary least squares (OLS) regression (stats v4.1.1 and olsrr v0.5.3). Analyses were stratified by nutrient treatment (organic or inorganic). We examined the following dependent variables: d values of plant shoot N and P, root N and P, NUE, PUE, shoot biomass, root biomass, RGR, and SRL. Independent predictors were fungal growth rate on three substrates (minimal media with organic N or organic P, and standard minimal media), NAG and AP enzyme activity, colonization rate, and habitat traits from the fungal site of origin (soil TIN, P, and HM). All predictor variables were centered and scaled (mean = 0, SD = 1) prior to analysis. Regression models include only 37-38 fungi because we could only successfully measure enzyme activity on 38 of the isolates and an additional fungus was missing NUE data from the organic nutrient treatment models. The data were averaged by fungal treatment within each nutrient condition to avoid pseudoreplication. Lastly, we used one-way ANOVAs to determine whether fungal colonization strategy (internal or external) influenced plant responses. To correct for multiple comparisons (n=20), we applied a Bonferroni-adjusted α = 0.0025 to the regressions and ANOVA.

## RESULTS

### Plant nitrogen and phosphorus

Shoot N and P content, which reflect plant nutrient acquisition unconfounded by fungal root colonization, depended on the interaction of fungus and nutrient treatment (*P* < 0.001; Table S3; Figures 2 A, B, S2). Shoot N exceeded uninoculated controls by 0.021-0.195 %N in 46% of fungal treatments in organic nutrient conditions and by 0.023-0.128 %N in 39% of fungal treatments in inorganic nutrient conditions (Figure 2A). The remaining fungal treatments largely had no effect on shoot N, with fewer than 8% of fungi in either nutrient treatment associated with lower shoot N (-0.024 to -0.058 %N) compared to controls. Shoot P was improved by 32% of fungi in the organic nutrient treatment (0.006-0.027 %P), compared to only 10% of fungi in the inorganic nutrient treatment (0.015-0.019 %P). More fungi also reduced shoot P relative to uninoculated controls in the organic (38%) compared to the inorganic (15%) treatment, and reductions were slightly larger in the latter (-0.007 to -0.018 %P) compared to the former (-0.004 to -0.012 %P). Shoot and root nutrients were generally poorly correlated (*r*^2^ = 0.056 to 0.161), except when plants were grown with organic P (*r*^2^ = 0.655).

**Figure 2.**
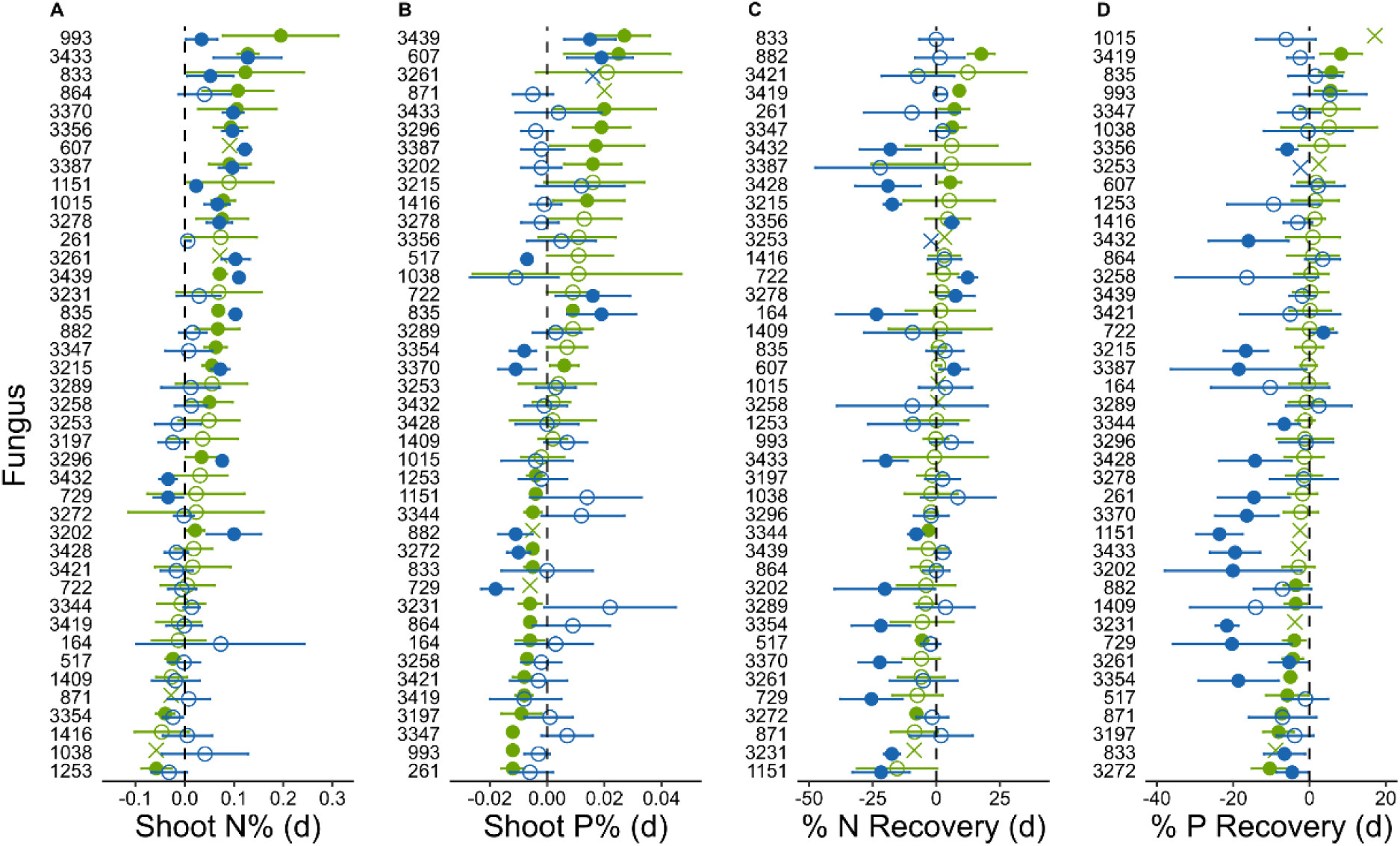
Plant nutrient responses to fungal isolates under inorganic (blue) and organic (green) nutrient treatments. Points show the mean effect size (d value) ± 95% confidence intervals. Solid circles indicate isolates whose 95% confidence interval do not overlap zero, open circles indicate intervals that overlap zero, and X symbols denote isolates with a single replicate (no confidence interval). The leading “NC0” has been removed from the fungal isolate names.

Fungal isolates differed substantially in their effects on plant nitrogen and phosphorus use efficiency, and these effects depended on nutrient source. Nitrogen use efficiency ranged from 6-85% and PUE ranged from 5-73% (Figure S2) across all treatments, reflecting significant fungus and nutrient treatment interactions (*P* < 0.001, Table S3). Relative to controls, there were five fungi that improved N recovery from organic nutrients (5.5-17.7% increase in %N) and four that improved N recovery from inorganic nutrients (3.1-12.3% increase in %N) (Figure 2C-D). Similarly, three fungi increased P recovery from organic sources (5.5-8.3 increase in %P) and one increased P recovery from inorganic sources (3.7%) relative to control plants. One fungus (NC00835) improved both N and P recovery in the organic nutrient treatment, while another (NC03387) improved both in the inorganic treatment. The majority of fungi had either no effect or a negative effect on plant NUE or PUE (Figure 2C-D). Negative effects were larger when inorganic nutrients were provided, with decreases of up to ∼25% relative to controls. In contrast, reductions in NUE and PUE in the organic nutrient treatments reached a maximum of ∼10%.

### Fungal trade-offs and specialization

When we examined the relationship between fungal effects on plant N and P content, we found trade-offs in nutrient acquisition, such that isolates associated with higher plant N% it tended to be associated with lower plant P% and vice versa, in both shoots and roots (Figure 3, S4, Table S4). In shoots, this trade-off only existed when nutrient sources were organic (*R*^2^ = 0.438, *P* < 0.001). In roots, there was a trade-off between N and P regardless of nutrient treatment and the slope of this relationship did not differ significantly between treatments (*R*^2^_organic_ = 0.575, *P*_organic_ < 0.001; *R*^2^_inorganic_ = 0.143, *P*_inorganic_ = 0.015).

**Figure 3.**
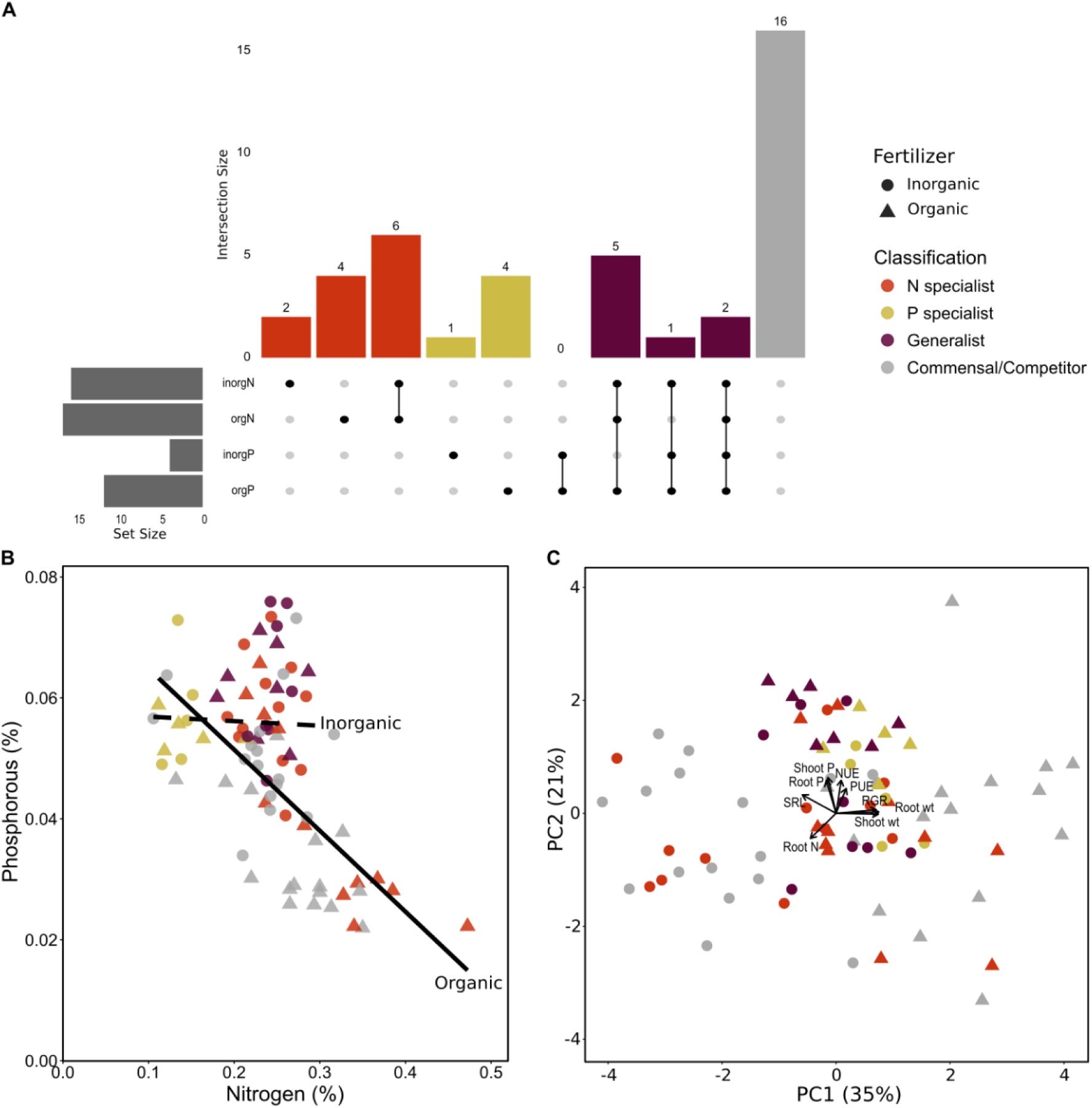
(A) Specialist and generalist fungal isolates based on significantly increased leaf nutrients across nutrient treatments. Each column represents a unique combination of nutrient benefits, with filled dots indicating the treatment conditions included in each combination. Bars are organized from specialist to generalist both within each nutrient (N or P) and across nutrients (N, P, both). Bar heights show the number of fungi in each intersection, while the set size bars show the number of fungi associated with benefits in each nutrient treatment. The final bar represents fungi that had only neutral or negative effects on plant nutrients (i.e., commensals and competitors). (B) Results of regressing plant phosphorous on nitrogen to determine whether there is a trade-off in resource acquisition in shoots where each point represents the average plant N% and P% for each fungus. Solid lines indicate significant relationships, while dashed lines are not significant. (C) Results of principle coordinate analysis of centered and standardized plant outcomes for the fungal isolates. In all plots, color indicates the classification of each fungal isolate as an N specialist (orange), P specialist (yellow), generalist (maroon), or those with neutral or negative effects (gray), while circles represent data from the inorganic nutrient treatment and triangles represent the organic nutrient treatment.

We further classified fungi as specialists or generalists based on their nutrient trade-offs and shared effects. Several fungi were N specialists that were beneficial only for plant N (NC00833, NC00993, NC01015, NC03215, NC03278, NC03356) or only for plants grown in either inorganic N (NC01151, NC03261) or organic N (NC00864, NC00882, NC03258, NC03347) (Figure 3, Table S1). Similarly, a small number of fungi specialized on organic P (NC00517, NC01416, NC03289, NC03354) or inorganic P (NC00722). Only two isolates, NC00835 and NC03439, were generalists that significantly improved both shoot N and P across the inorganic and organic nutrient treatments, but several other strains were beneficial for both shoot N and P in at least one nutrient treatment (NC00607, NC03202, NC03296, NC03370, NC03387, NC03433). The remaining strains were classified as commensals or competitors based on either no effect or reduced shoot nutrients, respectively, across the treatments (Figure 3, Table S1).

### Fungal traits as predictors

Although we observed substantial variation in both fungal traits and habitat traits (Figure S5), none were predictive of fungal effects on plant nutrient or growth outcomes (Table S5). However, there were two plant responses, shoot N% and root P%, in the inorganic nutrient treatment that could be predicted by this set of fungal and habitat traits when considering a less conservative alpha of 0.05 (Table S5).

## DISCUSSION

Root fungal endophytes generated pronounced variation in plant nutrient phenotypes, but whether fungi were beneficial or detrimental largely depended on the form of nutrients available. We were particularly interested in how these fungi affect plant access to organic nutrients, which are generally not bioavailable for plants. When only organic sources of N and P were available, approximately half of the 41 isolates increased shoot N and one third increased shoot P. Beneficial fungi improved shoot nutrients by an average of 0.08% N and 0.016% P relative to controls, but maximum benefits were ∼1.5× higher in the organic nutrient treatments than in the inorganic. These are modest but meaningful shifts relative to the observed ranges of shoot N (0.05-0.63%) and shoot P (0.02-0.09%). However, not all fungal treatments were beneficial and differences among fungal isolates in their effects on host nutrient acquisition under organic versus inorganic nutrient conditions, as well as apparent trade-offs in N and P, imply that root endophytic fungi vary in their capacity to acquire or mobilize distinct nutrient forms. These findings are consistent with growing evidence that endophyte-mediated benefits are conditional and can shift along mutualism–parasitism gradients depending on resource availability and environmental stressors (e.g.,(Kogel *et al*., 2006; Singh *et al*., 2011), which can challenge our ability to predict the outcomes of plant-fungal interactions.

Increases in plant nutrient content associated with the fungal treatments translated into nutrient recovery from organic sources of up to 47% for NUE and 35% for PUE, which were improvements of 20% and 17% relative to controls, respectively. The top organic nutrient recovery rates we observed exceed the 10-30% NUE and PUE previously reported for switchgrass (Owens *et al*., 2013; Tolofari *et al*., 2021; Osei *et al*., 2026) and are near the upper ranges reported for major cereal crops such as maize, wheat, and rice (Yu *et al*., 2022). Although direct comparisons across studies are complicated by differences in scale, nutrient sources, and nutrient loss pathways, this suggests that fungal symbionts can substantially enhance plant nutrient capture. Additional work is needed to understand if endophytic fungal associations with high organic matter in natural ecosystems (Kauppinen *et al*., 2014; Lee & Hawkes, 2021b) translate into enhanced abundance with manure or compost addition in agroecosystems, or whether other efforts—such as inoculations—would be required to support their role in fungal-mediated plant nutrient access.

The presence of organic and inorganic N specialists suggests a spectrum of N acquisition traits among the fungi that may also be influenced by host interactions. A subset of 12 phylogenetically diverse fungi comprised of 11 genera in 5 orders were N specialists on organic N, inorganic N, or both in terms of their benefits for the plant. Plants associated with the N specialist fungi also tracked the overall pattern of N:P tradeoffs from organic sources, but not inorganic. Complex organic sources of N are generally not preferred substrates for fungi given the need to invest in production of targeted enzymes and transporters (Kerkaert & Huberman, 2023). For example, one of our N specialists was putatively identified as a *Purpureocillium* sp., a genus that includes plant-beneficial taxa associated with enhanced nutrient acquisition and access to organic N pools through nematode parasitism and decomposition activity (Tang *et al*., 2023; Rigobelo *et al*., 2024). Similar to the functional diversity found here, studies of fungal litter decomposition indicate substantial variation among Ascomycetes in their preferences for N-, P-, or lignin-rich substrates (Manici *et al*., 2024). A comparable trait spectrum has been proposed for ectomycorrhizal fungi, with taxa ranging from “absorbers” to “miners” (Jörgensen *et al*., 2025). Although we observed both N generalists and specialists from the perspective of host N acquisition, substantial additional work on the endophytic fungi would be needed to identify relevant traits relating to variation in fungal ability to access or transform different forms of N or trade with plant hosts.

Only four endophytic fungi specialized on organic P, and all were at the low end for plant uptake of organic N consistent with a tradeoff for that specialization. The organic P specialists were putatively identified as a mycoparasite *Clonostachys* sp. (NC01416, (Sun *et al*., 2020), a potentially saprobic *Parastenospora* sp. (NC03354, (Crous *et al*., 2022), as well as strains of *Pezicula* (NC03289) and *Fonsecaea* (NC00517) where congeners are known to produce phosphatases (Kneipp *et al*., 2003; Luo *et al*., 2023). In contrast, specialization on inorganic P was limited to a single fungus (*Talaromyces* sp.) and no fungi were specialized on both forms. Moreover, negative effects were most frequently observed for shoot P and primarily in the organic nutrient treatment, where seven fungi were considered competitive including common strains of *Nigrospora* (NC03231) and *Penicillium* (NC00261). Fungi and plants were likely P-limited in our experiment given the application of only ∼2 mg/kg P, which could have heightened competition, particularly for taxa that efficiently mobilize limited P from organic matter but do not participate in trade with the plant (Johnson, 2010). Acquisition of organic P requires investment in extracellular phosphatases, and physical proximity to the site of mineralization may give fungi an advantage in acquisition. These patterns show that fungal effects on plant P were highly specialized, with organic P conditions promoting both competitive and a limited number of beneficial interactions.

Despite these specialized responses, a small subset of fungi functioned more broadly across nutrient forms as generalists. Just two isolates, both Dothideomycetes in the genus *Edenia* (NC00835, NC03439), significantly improved both shoot N and P across the inorganic and organic nutrient treatments, although several other strains trended in this direction. *Edenia* sp. are relatively common endophytic fungi, with some strains that are positively correlated with soil organic matter (Huang *et al*., 2026) or that enhance plant tolerance to drought stress (Li *et al*., 2021), but we found no prior reports on plant nutrient acquisition. Other fungi identified as generalists, including dark septate root endophytes, can decompose diverse organic substrates, increase plant access to micronutrients, promote plant growth, and contribute to rhizosphere network stability. This trend also aligns with the observed tradeoff between shoot N and P responses, with isolates that increased both nutrients relative to control plants tending to fall near the center of the N:P distribution. Whether nutrient generalists will make better biological inoculants than specialists will depend on needs as well as on their effects on other key plant traits such as productivity.

Although several fungi that reduced or increased shoot nutrients had the same effect on biomass, these were rarely aligned, suggesting variation in the balance between benefits and costs of fungal associations. Only two fungi, both organic P specialists, consistently improved plant biomass. Most of the fungi had no effect on plant biomass. This was particularly true in organic nutrient treatments, where 70-80% of fungi were neutral for both shoot and root biomass. In the inorganic treatments, more fungi resulted in reduced plant biomass, especially in roots (54%), although without any systematic changes in SRL or root-to-shoot ratio that would indicate shifts in plant investment. Reduced shoot biomass despite increased fungal-mediated tissue nutrients may indicate carbon costs of maintaining fungal symbionts. Although plant size was not correlated with fungal root colonization, a majority (∼60%) of fungi were found inside roots in our trait assay, allowing for the possibility of plant-fungal trade. Stable isotope tracer studies have demonstrated that many plant-fungal interactions involve reciprocal exchange of plant-derived carbon for fungal-acquired nutrients, with hosts allocating more carbon to fungal partners that deliver more limiting resources (e.g., Usuki & Narisawa, 2007; Vergara *et al*., 2017; Hoysted *et al*., 2023). Although carbon transfer was not measured here, variation in shoot biomass responses may reflect differences among fungal isolates in nutrient provisioning and carbon demand.

Variation in plant responses across fungal isolates and nutrient treatments suggests functional diversity among endophytes in nutrient degradation, acquisition, or transfer to the host plant. Many other root-associated fungi have been shown to have saprotrophic capabilities, including the enzymatic breakdown of complex organic substrates, which may enhance plant access to nutrients in organic forms (Knapp & Kovács, 2016). Although the general hydrolytic enzyme activities that we measured were not predictive of outcomes, this may reflect a mismatch between the enzymes measured and the specific enzymes required to access different nutrient pools. For example, casein breakdown requires proteases to break peptide bonds, which we did not measure. Similarly, phytic acid degradation is largely carried out by phytases, a subclass of phosphatase enzymes. In addition, fungal acid phosphatases are generally most active at very low pH of 2-5 (Ullah & Phillippy, 1994; Brazhnikova *et al*., 2022) and our assays at pH ≥ 5 likely did not represent in situ conditions where fungal organic acid production can significantly reduce local pH. The lack of predictive power found here might be improved by more targeted measurements aligned with specific nutrient acquisition pathways.

Functional trait frameworks are widely used to predict ecological roles and interaction outcomes, including in plant–fungal systems (Crowther *et al*., 2014; Zanne *et al*., 2020). Studies such as Giauque *et al*. (2019) or Khuna *et al*. (2021) have successfully linked fungal functional (e.g., osmotic sensitivity or mineral-solubilization) and habitat origin traits (e.g., climate) to plant growth and survival. However, in our study, fungal traits independent of the plant were not predictive of plant nutrient or growth responses. Functional traits measured in culture may not reflect fungal activity in the rhizosphere, as microbial nutrient-mobilizing abilities observed in laboratory assays do not necessarily scale to behavior in complex communities (Sharma *et al*., 2013). Even fungal colonization of the plant root was uncorrelated with host outcomes. Although root colonization is commonly measured for mycorrhizal fungi, it is not a universally effective predictor of fungal effects on plant resource acquisition or growth (Frew, 2025). Predicting plant-fungal interaction outcomes may require moving beyond general ecological or habitat traits towards more mechanistic or genetic traits that capture the potential capabilities of fungal nutrient processing or exchange.

Our experiment was conducted under controlled conditions where plants were paired with single fungal strains, which allowed us to isolate fungal effects on plant nutrients and growth. However, fungal behavior in natural systems reflects interactions among multiple microbial taxa, resource competition, and environmental heterogeneity (e.g., Frey-Klett *et al*., 2011; Marie Booth *et al*., 2019) that can shift observed outcomes, although we would still expect to see a range of fungal effects on the host. Extending this work to multi-isolate communities, field conditions, and additional functional trait assays will be important for evaluating the ecological generalizability of our findings.

These findings go beyond prior work focused on model taxa and indicate a general role for fungal root endophytes in plant nutrition that deserves further scrutiny, particularly to determine the mechanisms that support mycorrhiza-like behavior. The range of responses also underscores both the promise and the complexity of using root endophytes to enhance nutrient acquisition in low-input systems. Current agricultural practices rely on excessive applications of synthetic fertilizers and limiting both the economic and environmental burdens of these practices will be key to sustainable production. Based on our results, fungal endophytes that improve nutrient access could reduce reliance on synthetic fertilizers, but their nutrient-related context dependency indicates that this will not necessarily be straightforward. Future work integrating functional assays and genomic profiling of isolates, along with field-based validation, will be critical for discovering endophytes that enhance plant nutrient acquisition in practical ways.

## ACKNOWLEDGEMENTS

For assistance with laboratory and field work, we thank K. Harrell, C. Alexander, S. Merkel, H. Ervin, and X. Allen. Site access was generously provided by NCSU, North Carolina Department of Agriculture and Consumer Services, BASF Corp., Weyerhaeuser Co., C. Wilson, and E. Deal. This work was supported by a DOE Genomic Sciences Program SFA (SCW1039) subaward from Lawrence Livermore National Lab, by the Research Capacity Fund (HATCH), project award no. 7005451, from the U.S. Department of Agriculture’s National Institute of Food and Agriculture, and by funding from North Carolina State University to CVH. RH was supported by the Foundation for Food & Agriculture Research FFAR Fellows Program RDS-0000000002.

## AUTHOR CONTRIBUTIONS

The experiment was designed by CVH and MRL. The fungal culture collection was developed by NY. The experiment was carried out by MRL, NY, RH, and CVH. RH generated all nutrient data. Root-fungal microscopy images were generated by MK. Root lengths were obtained by MW. Root colonization was quantified by NL. RH analyzed data and RH and CVH wrote the first draft of the manuscript. All co-authors edited the manuscript.

## DATA and CODE AVAILABILITY

Isolate sequences will be available on the NCBI GenBank database. Data and R code will be available on GitHub.

