## Supplementary Material for "Diverse root fungal endophytes mediate plant access to soil nutrients"


| Item | Description | Page |
|---|---|---|
| | Supplemental Methods and Results | 2 |
| Figures | | |
| Fig. S1 | Phylogenetic autocorrelation heatmap | 3 |
| Fig. S2 | Plant nutrient responses to fungal and nutrient treatments | 4 |
| Fig. S3 | Plant growth responses to fungal and nutrient treatments | 5 |
| Fig. S4 | Fungal trade-offs in N and P acquisition | 6 |
| Fig. S5 | Fungal traits | 7 |
| Fig. S6 | Micrographs – external colonization | 8 |
| Tables | | |
| Table S1 | Taxonomic and phylogenetic identification of isolates | 10 |
| Table S2 | Results of testing for phylogenetic autocorrelation | 11 |
| Table S3 | Results of ANOVAs of plant response regressed on treatment | 12 |
| Table S4 | Results of linear regression to test for N:P tradeoff | 13 |
| Table S5 | Results of OLS regression to predict outcomes from traits | 14 |
| Table S6 | Results of ANOVA to predict outcomes from colonization strategy | 15 |

### Supplemental Methods and Results

#### Fungal imaging

To ensure all fungi were colonizing the root after storage, we assayed fungal root colonization in switchgrass. For this purpose, we grew sterile switchgrass seedlings inoculated with each fungus for 4 weeks as described earlier. Plants were harvested and roots were fixed in 50% ethanol before being stained with WGA-AlexaFluor488. They were imaged using both Differential Interference Contrast (DIC) and fluorescence microscopy with the resulting images overlayed to visualize the spatial orientation of the fungus within the root. Fungal colonization was confirmed for all taxa (Figure 1, S5).

#### Root nutrients

Belowground, root P was also affected by the interaction of fungus and nutrient treatments ($P <$ 0.001; Table S3; Figure 2D). Most fungi did not affect root P (51% organic, 83% inorganic). In the organic nutrient treatment, root P was improved by up to 0.01 %P in 16% of fungi and reduced by a maximum of 0.003 %P in 32% of fungi relative to uninoculated controls. The effects of fungi on root P in organic conditions were consistent with shoot P in 83% of cases with negative impacts and in 67% of cases with positive impacts. In the inorganic nutrient treatment, increased root P (0.015-0.019 %P) was associated with only 13% of fungi and decreased root P (-0.007 to -0.018 %P) was found with only 5% of fungi compared to controls, and these were inconsistent with shoot P effects.

Root N was only significantly affected by the nutrient treatments ($P$ = 0.002, Table S3, Figure 2B), which were on average more negative in the organic (-0.024 ± 0.005 SE) compared to the inorganic (-0.001 ± 0.002 SE) nutrient treatment compared to uninoculated controls. There was also a trend in the fungus by nutrient treatment interaction ($P$ = 0.053; Table S3). In planned comparisons, most fungi had no significant effect on root N (49% organic, 71% inorganic) compared to uninoculated controls and very few fungi improved root N in either nutrient treatment (0% organic, 10% inorganic), the latter of which were uncorrelated with shoot N (Figure 2B). Among fungal treatments that caused a decrease in root N relative to controls (49% organic, 20% inorganic), ~50% of cases were associated with significant increases in shoot N.

#### Plant growth and size

Shoot dry weight ranged from 0.01-2.09 g, root dry weight ranged from 0.01-3.46 g, and we observed 7- to 12-fold variation in RGR (excluding one sample that showed slightly negative growth) (Figure S3). Plant shoot biomass, root biomass, and RGR were also affected by the interaction of fungus and nutrient treatment (P < 0.001; Table S3). Fungal effects on plant growth were predominantly negative (Figure S3). In the organic nutrient treatment, 17% of fungi reduced shoot biomass (-0.15 to -0.40g), 20% reduced root biomass (-0.23 to -0.45g), and 12% reduced RGR (-0.08 to -0.13 cm/day) compared to controls. In the inorganic nutrient treatment, these proportions increased to 34% for shoot biomass (-0.25 to -0.82g), 54% for root biomass (-0.30 to -1.16g), and 24% for RGR (-0.06 to -0.24 cm/day). Only a small number of fungi (n=1-6) significantly increased either shoot biomass (by ≤ 0.32g), root biomass (0.23g) or RGR (by ≤ 0.14 cm/day) in one of the nutrient treatments compared to controls. SRL (d) was unaffected by either fungus or nutrient treatment (P > 0.006, Table S3), despite up to 97-fold variation across the treatments.

**Figure S1**. Trait heat map. Plant response traits were shoot weight, root weight, relative height growth rate (RGR), shoot %N, shoot %P, root %N, root %P, nitrogen use efficiency (NUE), and phosphorus use efficiency (PUE). All plant response traits are shown for both the organic (o) and inorganic (i) treatments. Fungal traits were % root colonization, growth rates (GR) in culture with M9 minimal media (MM), with MM where casein replaced inorganic N (GR+C), and with MM where phytic acid replaced inorganic P (GR+P), and activities of acid phosphatase (AP) and N-acetyl glucosaminidase (NAG) enzymes. A subset of the fungal isolates were missing data for NUE or enzymes, which are denoted by white boxes with a slash. Traits are in columns and fungi are in rows. The final column indicates whether we classified a fungus as a generalist or specialist (see Fig. 3 legend). Fungal rows are arranged by phylogeny, with the colored box representing class.

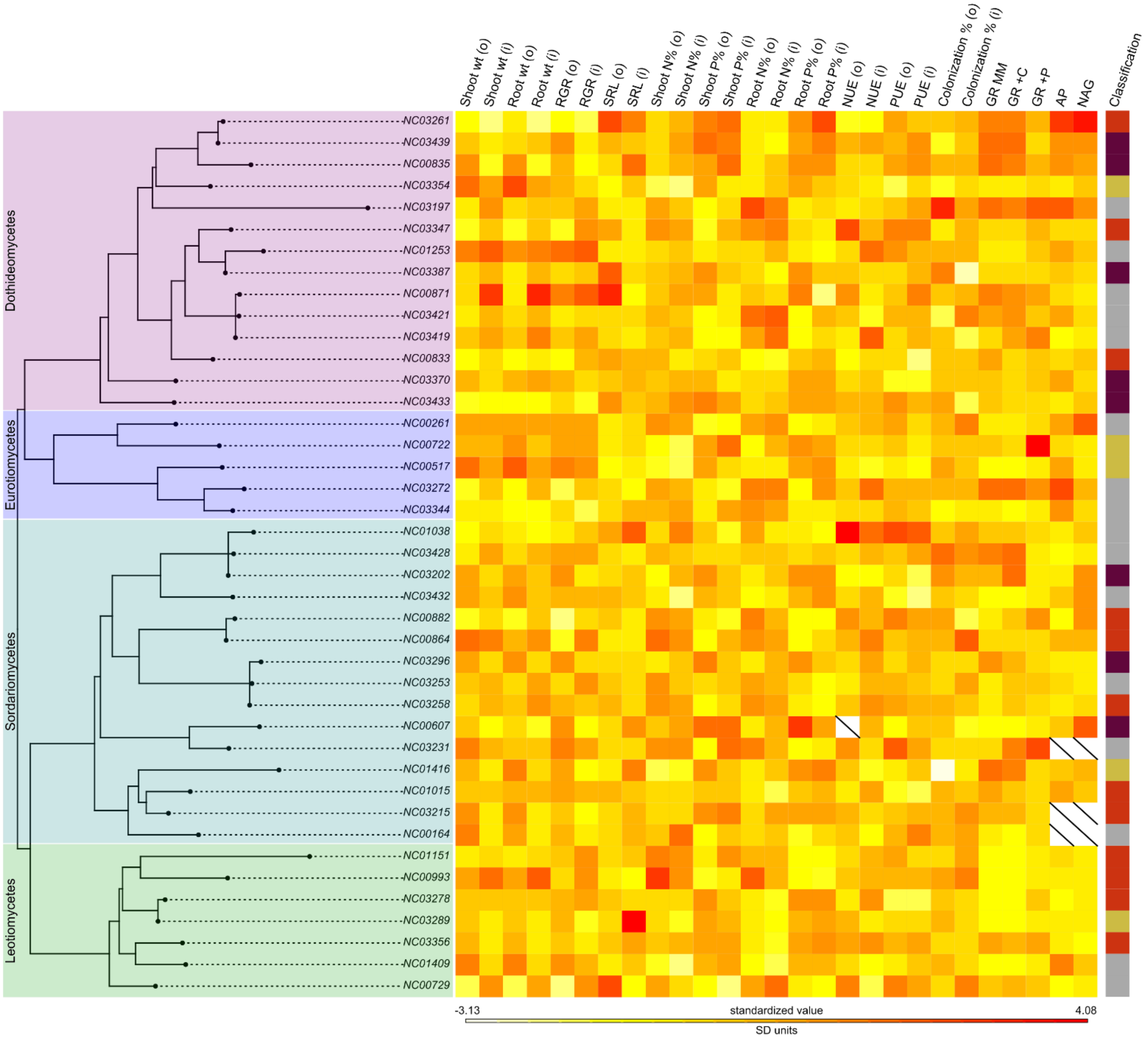

**Figure S2**. Plant nutrient responses to fungal isolates under inorganic (blue) and organic (green) nutrient treatments. In panels A-F, points are mean ± 1 SD. Isolates with a single replicate for that measurement are indicated by “×”, while points in panels G-H show the mean effect size (d value) ± 95% confidence intervals, and solid circles indicate isolates whose 95% confidence interval do not overlap zero and open circles indicate intervals that overlap zero. The leading “NC0” has been removed from the fungal isolate names for space. Fungi are arranged by descending numeric order within each temporal batch, separated by the gray horizontal line.

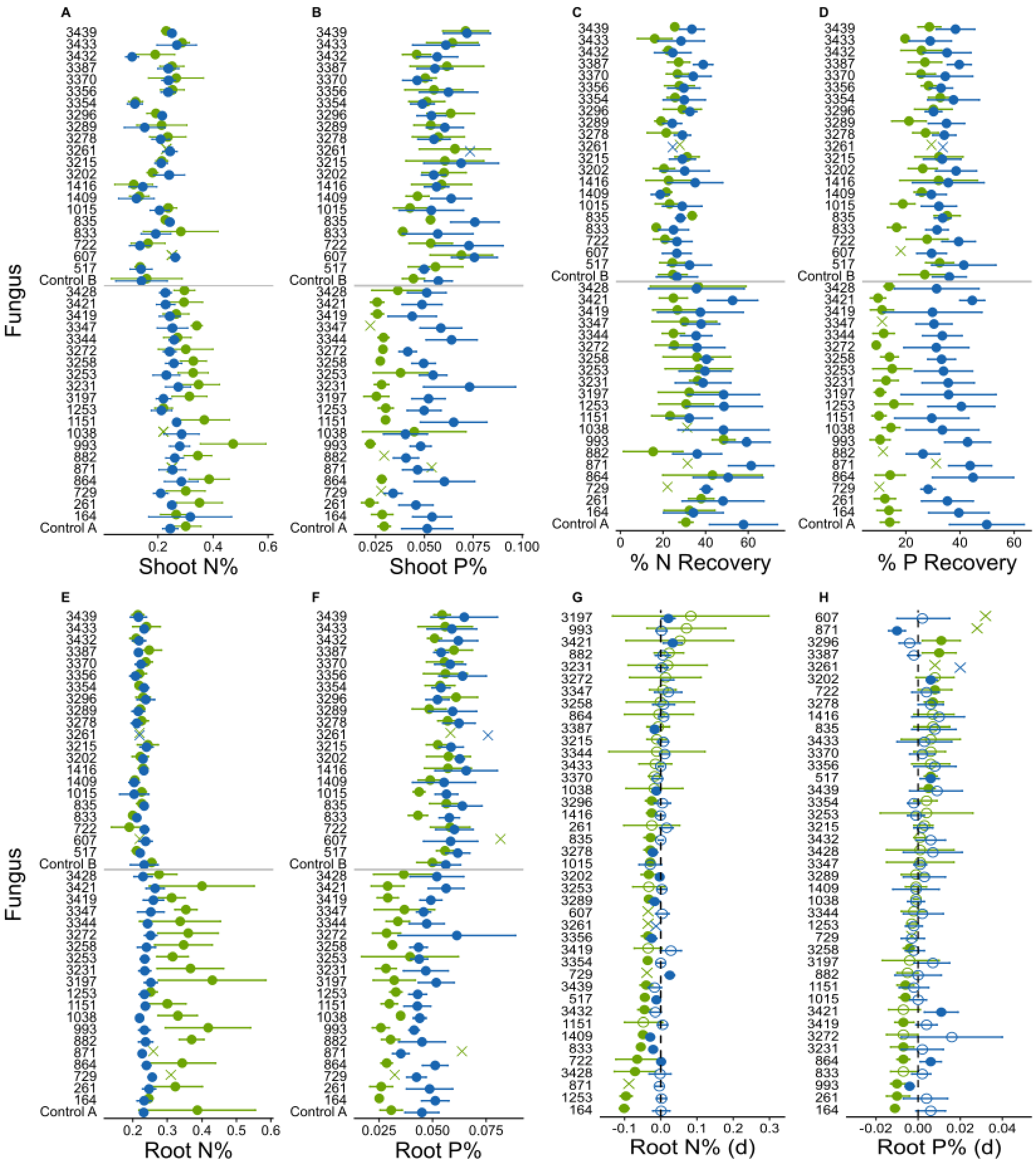

**Figure S3**. Plant growth responses to fungal isolates under inorganic (blue) and organic (green) nutrient treatments. In the top row, points are mean ± 1 SD. In the second row points show the mean effect size (d value) ± 95% confidence intervals, and solid circles indicate isolates whose 95% confidence interval do not overlap zero and open circles indicate intervals that overlap zero. Fungi are either arranged by descending numeric order within each temporal batch (top) or ordered by performance on the organic nutrient treatment (bottom). The leading “NC0” has been removed from fungal isolate names.

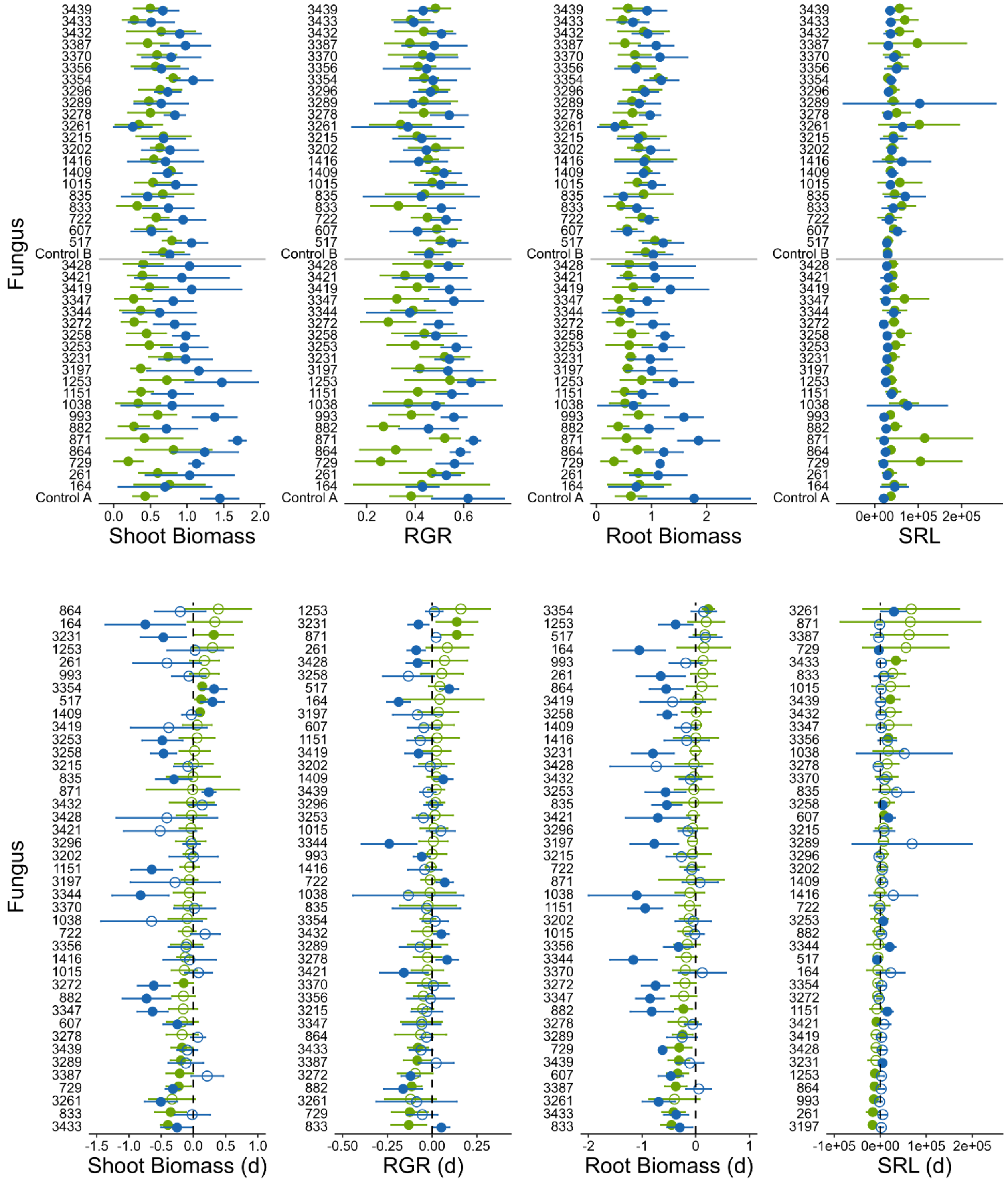

**Figure S4**. Results of regressing plant phosphorous on nitrogen to determine whether there is a trade-off in resource acquisition in roots.

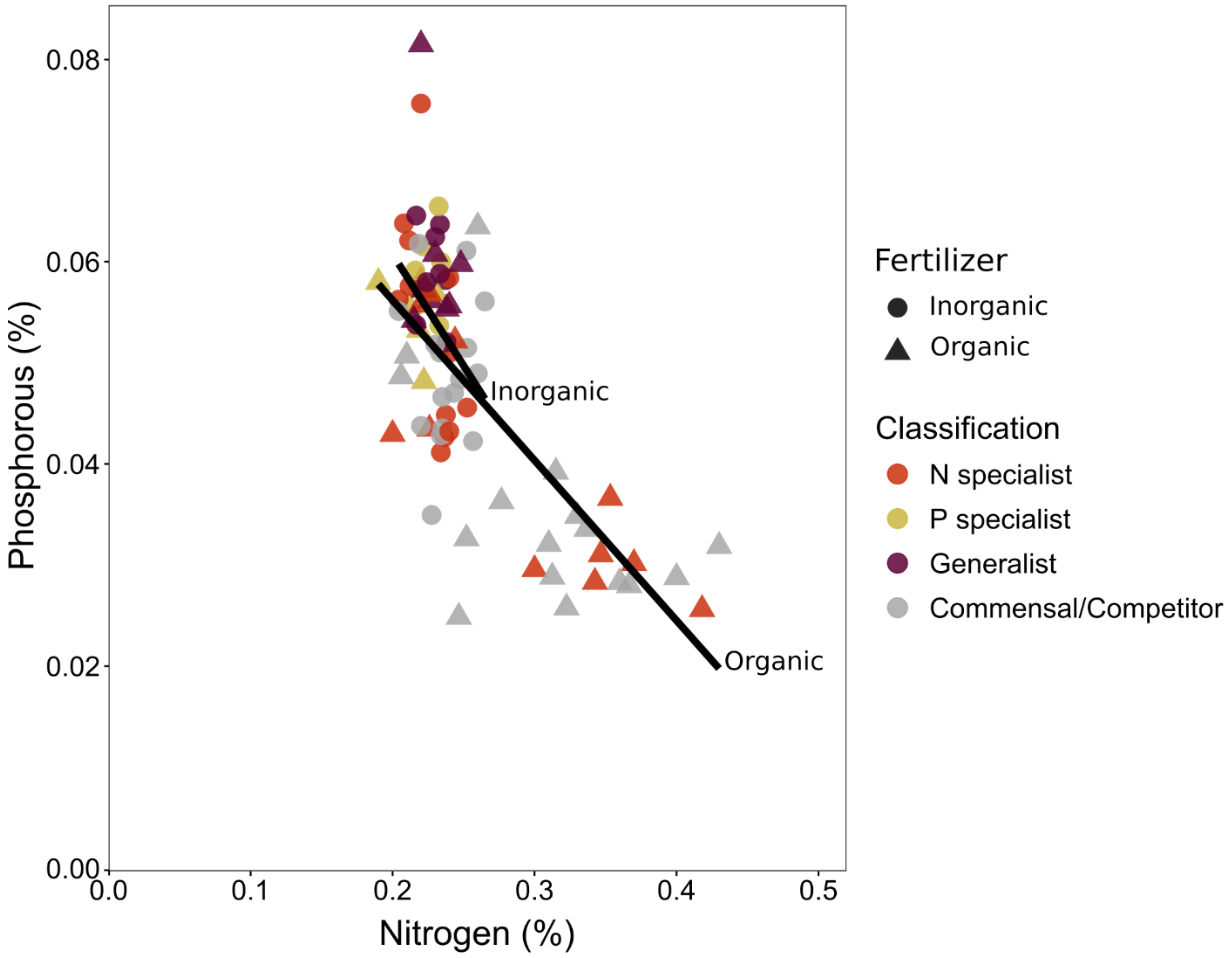

**Figure S5.** Fungal traits measured in plants of (A, B) colonization, or in culture of (C, D, E) growth rate and (F, G) enzyme activity (missing 3 fungi), and characteristics of the (H, I, J) habitats from which fungi were isolated. Bars show the mean ± 1 SD.

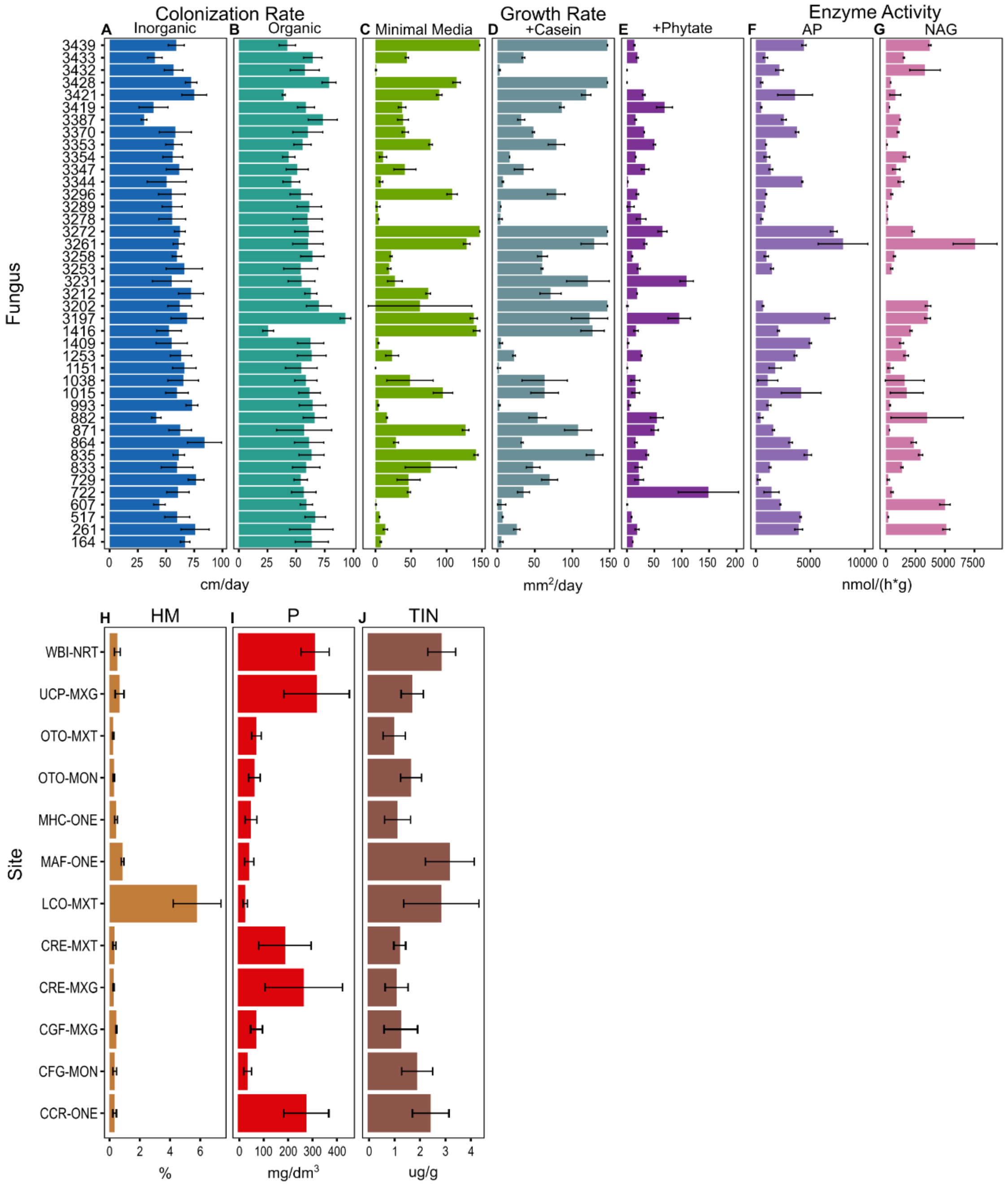

**Figure S6**. Composite images of fluorescence and DIC microscopy of roots where the fungi fluoresce green. Scale bar represents 50 µm.

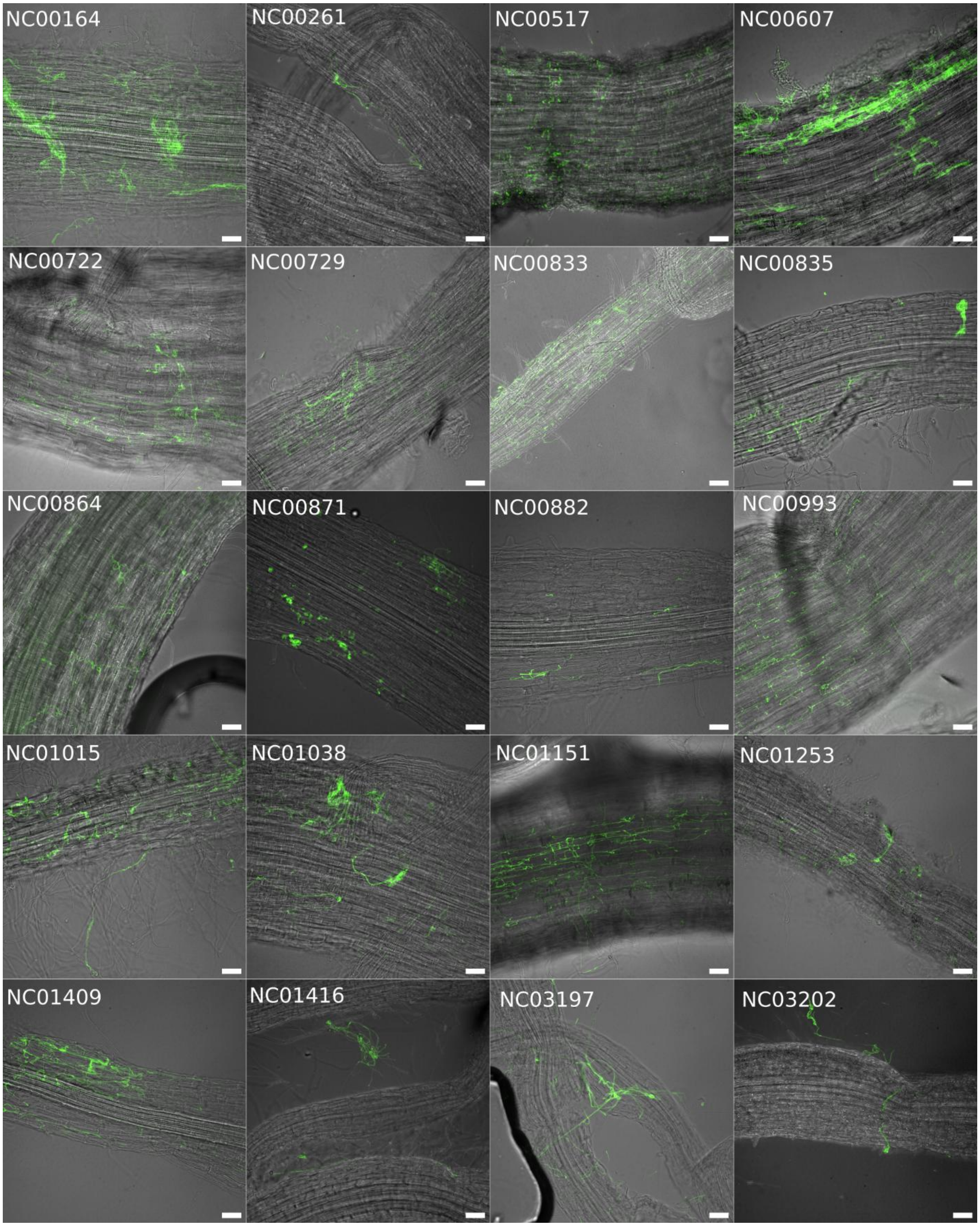

**Figure S6** (continued)

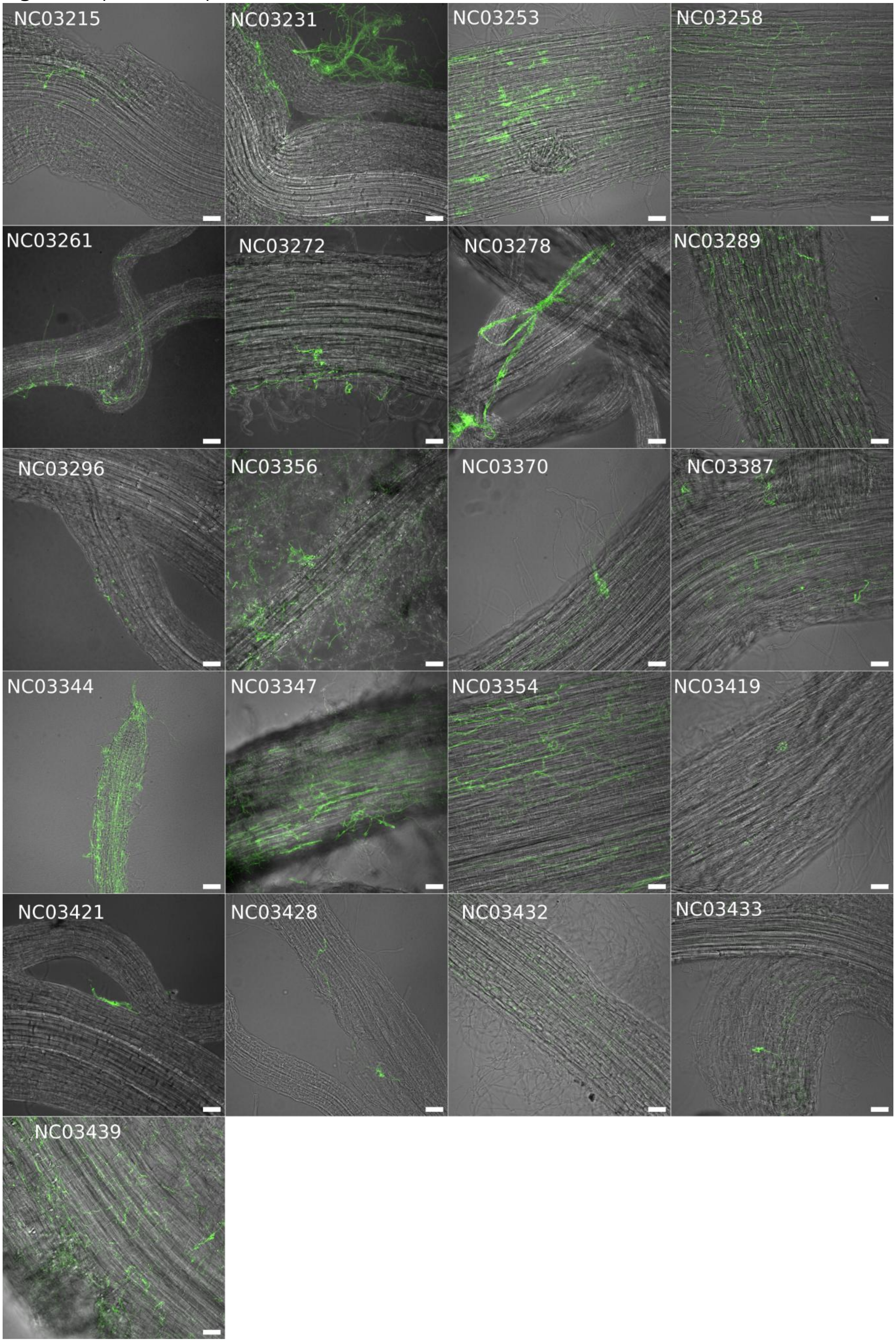

**Table S1**. Identification of the fungal isolates based on placing the LSU rDNA or LSU and ITS rDNA sequences (for a subset of isolates with successful ITS sequencing, n = 38) on a T-BAS reference tree. Classification based on mean effect size for shoot nutrients under either nutrient condition (organic = o, inorganic = i). Most isolate ITS sequences are the full sequence, except NC03433 is ITS2 only.

| Isolate | Site | Isolate LSU Accession | Isolate ITS Accession | T-BAS Identification | Classification |
|---|---|---|---|---|---|
| NC00164 | CRE-MXT | PZ441615 | | Trichoderma | Competitor P (o) |
| NC00261 | OTO-MXT | PZ441616 | PZ435688 | Penicillium | Competitor P (o) |
| NC00517 | WBI-NRT | PZ441617 | | Fonsecaea | Specialist P (o) |
| NC00607 | MAF-ONE | PZ441618 | PZ435689 | Allocryptovalsa | Generalist |
| NC00722 | OTO-MON | PZ441619 | PZ435690 | Talaromyces | Specialist P (i) |
| NC00729 | OTO-MON | PZ441620 | PZ435691 | Pseudeurotium | Competitor (i) |
| NC00833 | CRE-MXT | PZ441621 | PZ435692 | Pseudoxylomyces | Specialist N |
| NC00835 | CRE-MXT | PZ441622 | PZ435693 | Edenia | Generalist |
| NC00864 | UCP-MXG | PZ441623 | | Arcopilus | Specialist N (o) |
| NC00871 | CRE-MXT | PZ441624 | PZ435694 | Flavomyces | Commensal |
| NC00882 | LCO-MXT | PZ441625 | PZ435695 | Arcopilus | Specialist N (o) |
| NC00993 | UCP-MXG | PZ441626 | PZ435696 | Rommelaarsia | Specialist N |
| NC01015 | CRE-MXG | PZ441627 | PZ435697 | Purpureocillium | Specialist N |
| NC01038 | CRE-MXG | PZ441628 | PZ435698 | Magnaporthiopsis | Commensal |
| NC01151 | MHC-ONE | PZ441629 | PZ435699 | Mollisia | Specialist N (i) |
| NC01253 | CGF-MON | PZ441630 | PZ435700 | Poaceascoma | Competitor (o) |
| NC01409 | CGF-MXG | PZ441631 | | Gyoerffyella | Commensal |
| NC01416 | CGF-MXG | PZ441632 | PZ435701 | Clonostachys | Specialist P (o) |
| NC03197 | CCR-ONE | PZ441633 | PZ435702 | Didymella | Competitor P (o) |
| NC03202 | CCR-ONE | PZ441634 | PZ435703 | Magnaporthiopsis | Generalist |
| NC03215 | CCR-ONE | PZ441635 | PZ435704 | Pochonia | Specialist N |
| NC03231 | CCR-ONE | PZ441636 | PZ435705 | Nigrospora | Competitor P (o) |
| NC03253 | CCR-ONE | PZ441637 | PZ435706 | Codinaea | Commensal |
| NC03258 | CCR-ONE | PZ441638 | PZ435707 | Codinaea | Specialist N (o) |
| NC03261 | CCR-ONE | PZ441639 | PZ435708 | Edenia | Specialist N (i) |
| NC03272 | CCR-ONE | PZ441640 | PZ435709 | Metulocladosporiella | Competitor P |
| NC03278 | UCP-MXG | PZ441641 | PZ435710 | Pezicula | Specialist N |
| NC03289 | UCP-MXG | PZ441642 | PZ435711 | Pezicula | Specialist P (o) |
| NC03296 | UCP-MXG | PZ441643 | PZ435712 | Codinaea | Generalist |
| NC03344 | CRE-MXT | PZ441644 | PZ435713 | Metulocladosporiella | Competitor P (o) |
| NC03347 | CRE-MXT | PZ441645 | PZ435714 | Poaceascoma | Specialist N (o) |
| NC03354 | CRE-MXT | PZ441646 | PZ435715 | Parastenospora | Specialist P (o) |
| NC03356 | CRE-MXT | PZ441647 | PZ435716 | Collembolispora | Specialist N |
| NC03370 | CRE-MXT | PZ441648 | | Tetraploa | Generalist |
| NC03387 | CRE-MXT | PZ441649 | PZ435717 | Poaceascoma | Generalist |
| NC03419 | CGF-MXG | PZ441650 | PZ435718 | Flavomyces | Competitor P (o) |
| NC03421 | CGF-MXG | PZ441651 | PZ435719 | Flavomyces | Competitor P (o) |
| NC03428 | CGF-MXG | PZ441652 | PZ435720 | Magnaporthiopsis | Commensal |
| NC03432 | CGF-MXG | PZ441653 | PZ435721 | Ophioceras | Competitor N (i) |
| NC03433 | CGF-MXG | PZ441654 | PZ436069 | Lindgomyces | Generalist |
| NC03439 | CGF-MXG | PZ441655 | PZ435722 | Edenia | Generalist |

**Table S2**. Results of using phylogenetic autocorrelation to determine whether fungal relatedness constrained fungal treatment effects on plant nutrient acquisition and growth in each nutrient treatment. A subset of the 41 fungal isolates had enzyme activity data (n=38) and available data for calculating NUE in the organic treatment (n=40).

| Test | Obs | Std.Obs | Alter | *P* |
|---|---|---|---|---|
| Fungus | 0.917 | 9.023 | greater | 0.001 |
| *Plant effects* | | | | |
| SRL (organic) | -0.003 | -0.063 | greater | 0.460 |
| SRL (inorganic) | -0.018 | -0.150 | greater | 0.527 |
| Shoot biomass (organic) | 0.004 | 0.031 | greater | 0.466 |
| Shoot biomass (inorganic) | 0.015 | 0.151 | greater | 0.406 |
| Root biomass (organic) | 0.013 | 0.127 | greater | 0.429 |
| Root biomass (inorganic) | -0.005 | -0.038 | greater | 0.503 |
| RGR (organic) | 0.002 | 0.024 | greater | 0.466 |
| RGR (inorganic) | 0.045 | 0.455 | greater | 0.304 |
| Shoot P (inorganic) | 0.007 | 0.083 | greater | 0.429 |
| Shoot P (organic) | -0.046 | -0.419 | greater | 0.643 |
| Shoot N (inorganic) | -0.004 | -0.045 | greater | 0.492 |
| Shoot N (organic) | -0.056 | -0.532 | greater | 0.700 |
| Root P (inorganic) | 0.063 | 0.624 | greater | 0.259 |
| Root P (organic) | -0.043 | -0.417 | greater | 0.660 |
| Root N (inorganic) | -0.186 | -1.882 | greater | 0.986 |
| Root N (organic) | -0.070 | -0.703 | greater | 0.743 |
| PUE (inorganic) | -0.105 | -1.068 | greater | 0.860 |
| PUE (organic) | -0.023 | -0.241 | greater | 0.561 |
| NUE (inorganic) | 0.122 | 1.159 | greater | 0.127 |
| *Fungal traits* | | | | |
| Colonization rate (inorganic) | 0.070 | 0.646 | greater | 0.249 |
| Colonization rate (organic) | -0.026 | -0.267 | greater | 0.596 |
| Growth rate - casein | 0.004 | 0.005 | greater | 0.472 |
| Growth rate - phytate | -0.081 | -0.898 | greater | 0.813 |
| Growth rate - minimal media | -0.136 | -1.237 | greater | 0.890 |

| Test | Obs | Std.Obs | Alter | *P* |
|---|---|---|---|---|
| Fungus | 0.914 | 8.577 | greater | 0.001 |
| *Enzyme* | | | | |
| AP | -0.141 | -1.321 | greater | 0.921 |
| NAG | -0.002 | -0.007 | greater | 0.480 |

| Test | Obs | Std.Obs | Alter | *P* |
|---|---|---|---|---|
| Fungus | 0.055 | 0.483 | greater | 0.318 |
| *Plant effects* | | | | |
| NUE (organic) | -0.140 | -1.534 | greater | 0.966 |

**Table S3**. Results of ANOVAs of plant (a-d, i-j) nutrient and (e-h) growth outcomes on fungus and fertilizer treatments, using Bonferroni-adjusted alpha = 0.00625. Analysis of NUE included data for only 40 fungi, due to missing data in the organic nutrient treatment.

**a. Shoot N**

| | SS | df | *F* | *P* |
|---|---|---|---|---|
| (Intercept) | 0.016 | 1 | 5.823 | 0.016 |
| fungus | 0.477 | 40 | 4.379 | <0.001 |
| fert | 0.012 | 1 | 4.571 | 0.033 |
| fungus:fert | 0.205 | 40 | 1.885 | 0.002 |
| Residuals | 0.786 | 289 | | |

**b. Shoot P**

| | SS | df | *F* | *P* |
|---|---|---|---|---|
| (Intercept) | 1.25e-5 | 1 | 0.097 | 0.756 |
| fungus | 0.015 | 40 | 2.954 | <0.001 |
| fert | 9.40e-5 | 1 | 0.727 | 0.395 |
| fungus:fert | 0.012 | 40 | 2.409 | <0.001 |
| Residuals | 0.035 | 270 | | |

**c. Root N**

| | SS | df | *F* | *P* |
|---|---|---|---|---|
| (Intercept) | 5.33e-6 | 1 | 0.003 | 0.953 |
| fungus | 0.044 | 40 | 0.717 | 0.897 |
| fert | 0.016 | 1 | 10.106 | 0.002 |
| fungus:fert | 0.089 | 40 | 1.434 | 0.053 |
| Residuals | 0.397 | 256 | | |

**d. Root P**

| | SS | df | *F* | *P* |
|---|---|---|---|---|
| (Intercept) | 1.16e-4 | 1 | 1.464 | 0.227 |
| fungus | 0.005 | 40 | 1.443 | 0.050 |
| fert | 4.21e-4 | 1 | 5.304 | 0.022 |
| fungus:fert | 0.007 | 40 | 2.164 | <0.001 |
| Residuals | 0.020 | 258 | | |

**e. Shoot biomass**

| | SS | df | *F* | *P* |
|---|---|---|---|---|
| (Intercept) | 2.235 | 1 | 20.266 | <0.001 |
| fungus | 19.890 | 40 | 4.510 | <0.001 |
| fert | 2.591 | 1 | 23.498 | <0.001 |
| fungus:fert | 13.138 | 40 | 2.979 | <0.001 |
| Residuals | 38.703 | 351 | | |

**f. Root biomass**

| | SS | df | *F* | *P* |
|---|---|---|---|---|
| (Intercept) | 4.446 | 1 | 35.989 | <0.001 |
| fungus | 25.885 | 40 | 5.238 | <0.001 |
| fert | 3.228 | 1 | 26.128 | <0.001 |
| fungus:fert | 15.81 | 40 | 3.199 | <0.001 |
| Residuals | 43.121 | 349 | | |

**g. RGR**

| | SS | df | *F* | *P* |
|---|---|---|---|---|
| (Intercept) | 0.140 | 1 | 10.982 | 0.001 |
| fungus | 1.214 | 40 | 2.377 | <0.001 |
| fert | 0.118 | 1 | 9.252 | 0.003 |
| fungus:fert | 1.001 | 40 | 1.960 | 0.001 |
| Residuals | 4.481 | 351 | | |

**h. SRL**

| | SS | df | *F* | *P* |
|---|---|---|---|---|
| (Intercept) | 2e9 | 1 | 0.898 | 0.344 |
| fungus | 1e11 | 40 | 1.148 | 0.255 |
| fert | 2e9 | 1 | 0.742 | 0.390 |
| fungus:fert | 9e10 | 40 | 0.982 | 0.505 |
| Residuals | 8e11 | 351 | | |

**i. % N recovery**

| | SS | df | *F* | *P* |
|---|---|---|---|---|
| (Intercept) | 1659.5 | 1 | 16.460 | <0.001 |
| fungus | 19446.4 | 39 | 4.946 | <0.001 |
| fert | 951.1 | 1 | 9.433 | 0.002 |
| fungus:fert | 9213 | 39 | 2.343 | <0.001 |
| Residuals | 23189.5 | 230 | | |

**j. % P recovery**

| | SS | df | *F* | *P* |
|---|---|---|---|---|
| (Intercept) | 3133.9 | 1 | 49.699 | <0.001 |
| fungus | 3758.2 | 40 | 1.490 | 0.037 |
| fert | 800.8 | 1 | 12.699 | <0.001 |
| fungus:fert | 6407.7 | 40 | 2.540 | <0.001 |
| Residuals | 15196.8 | 241 | | |

**Table S4**. Results of linear regression of plant P on N from each of the nutrient treatments.

| Treatment | Compartment | β ± SE | *F* (df1, df2) | *P* | $R^2$ |
|---|---|---|---|---|---|
| Organic | Shoot | -0.13 ± 0.02 | 30.4 (1, 39) | <0.001 | 0.438 |
| | Root | -0.16 ± 0.02 | 52.8 (1, 39) | <0.001 | 0.575 |
| Inorganic | Shoot | -0.01 ± 0.03 | 0.1 (1, 39) | 0.823 | 0.001 |
| | Root | -0.22 ± 0.09 | 6.5 (1, 39) | 0.015 | 0.143 |

**Table S5**. Results of OLS regressions for predicting plant nutrient and growth outcomes from standardized fungal traits. Traits included colonization rate of the plants (colonization), growth rate on minimal media (GR) or minimal media with the N and P sources of casein (GR N) or phytate (GR P), enzyme activity (AP, NAG), and habitat of isolation characteristics of total inorganic nitrogen (TIN), phosphorus (P), and percent humic matter (HM). No coefficients were significant after Bonferroni correction (α = 0.003). Analyses included data for 38 fungi out of 41 total, except NUE in the organic nutrient treatment which included 37, due to missing data.

**a. Model-level summaries**

| Treatment | Outcome | *F* (df1, df2) | *P* | Adj. $R^2$ |
|---|---|---|---|---|
| Organic | Shoot N | 0.92 (9, 28) | 0.526 | -0.021 |
| | Shoot P | 1.47 (9, 28) | 0.208 | 0.102 |
| | Root N | 1.14 (9, 28) | 0.372 | 0.032 |
| | Root P | 1.81 (9, 28) | 0.111 | 0.164 |
| | Shoot biomass | 0.89 (9, 28) | 0.550 | -0.029 |
| | Root biomass | 0.58 (9, 28) | 0.804 | -0.115 |
| | RGR | 0.71 (9, 28) | 0.695 | -0.076 |
| | SRL | 0.71 (9, 29) | 0.695 | -0.076 |
| | NUE | 1.51 (9, 27) | 0.195 | 0.113 |
| | PUE | 0.51 (9, 28) | 0.854 | -0.135 |
| Inorganic | Shoot N | 2.54 (9, 28) | 0.029 | 0.272 |
| | Shoot P | 1.00 (9, 28) | 0.460 | 0.001 |
| | Root N | 1.55 (9, 28) | 0.181 | 0.117 |
| | Root P | 3.01 (9, 28) | 0.012 | 0.329 |
| | Shoot biomass | 1.11 (9, 28) | 0.391 | 0.025 |
| | Root biomass | 0.84 (9, 28) | 0.584 | -0.040 |
| | RGR | 1.28 (9, 28) | 0.291 | 0.064 |
| | SRL | 0.49 (9, 28) | 0.867 | -0.141 |
| | NUE | 1.86 (9, 28) | 0.100 | 0.174 |
| | PUE | 1.12 (9, 28) | 0.383 | 0.028 |

**Table S5. Continued**

**b. Standardized regression coefficients (β)**

| Treatment | Trait | Shoot N | Shoot P | Root N | Root P | Shoot biomass | Root biomass | RGR | SRL | NUE | PUE |
|---|---|---|---|---|---|---|---|---|---|---|---|
| Organic | *Plant* | | | | | | | | | | |
| | Colonization | 0.014 | -0.001 | -2.30e-4 | -0.001 | -1.65e-5 | -0.012 | 0.012 | 1281.117 | 0.983 | -0.943 |
| | *Culture* | | | | | | | | | | |
| | GR N | -0.034 | -0.008 | 0.014 | -0.008 | 0.102 | 0.099 | 0.008 | -12885.158 | 0.765 | 0.747 |
| | GR P | -0.007 | -0.002 | 0.004 | -4.04e-4 | 0.003 | 0.016 | -0.010 | -2302.032 | -0.347 | 0.465 |
| | GR | 0.028 | 0.011 | -0.015 | 0.010 | -0.161 | -0.144 | -0.007 | 18370.576 | -0.598 | 0.576 |
| | AP | -0.015 | -0.006 | 0.012 | -0.005 | 0.097 | 0.054 | 0.004 | -8568.738 | -0.054 | -0.107 |
| | NAG | 0.017 | 0.004 | -0.002 | 0.002 | -0.037 | -0.043 | -0.015 | 6689.392 | 1.031 | -0.287 |
| | *Habitat* | | | | | | | | | | |
| | TIN | -0.008 | 0.004 | -0.006 | 0.005 | -0.021 | -0.016 | 0.013 | -772.891 | -1.710 | -0.306 |
| | P | 0.013 | -2.75e-4 | 0.015 | -0.002 | 0.014 | 0.013 | -0.021 | 682.151 | 2.838 | 0.875 |
| | HM | 0.008 | -0.003 | 0.014 | -0.003 | -0.004 | -0.010 | -0.023 | -2863.875 | -1.359 | 0.104 |
| Inorganic | *Plant* | | | | | | | | | | |
| | Colonization | -0.022 | -0.001 | 0.004 | -2.64e-4 | -0.046 | -0.103 | 0.011 | -2604.891 | -2.995 | -2.085 |
| | *Culture* | | | | | | | | | | |
| | GR N | -0.011 | -0.005 | 0.010 | 0.004 | -0.257 | -0.172 | -0.078 | 1188.065 | -8.581 | -4.557 |
| | GR P | -0.009 | 2.82e-4 | 0.004 | 0.001 | 0.017 | -0.004 | 0.004 | -1427.487 | -0.663 | -0.064 |
| | GR | 0.023 | 0.004 | -0.010 | -0.002 | 0.238 | 0.177 | 0.069 | -1353.192 | 10.163 | 4.505 |
| | AP | -0.016 | -6.24e5 | 0.001 | 0.002 | -0.098 | -0.086 | -0.024 | -1230.613 | -3.755 | -1.281 |
| | NAG | 0.025 | 0.003 | -0.002 | 0.001 | 0.013 | -0.010 | 0.001 | 4960.563 | 3.586 | 2.236 |
| | *Habitat* | | | | | | | | | | |
| | TIN | -0.004 | 0.001 | 0.003 | -5.37e-5 | 0.047 | 0.041 | 0.013 | -3474.216 | -0.124 | 0.063 |
| | P | 0.016 | -0.002 | -0.004 | 2.59e-4 | 0.016 | 0.040 | -0.004 | 4209.088 | 2.590 | 1.765 |
| | HM | -0.005 | -0.003 | -0.001 | -4.34e-4 | -0.103 | -0.091 | -0.024 | 631.563 | -2.654 | -2.641 |
